# SARS-CoV-2 Nsp15 antagonizes the cGAS-STING-mediated antiviral innate immune responses

**DOI:** 10.1101/2024.09.05.611469

**Authors:** Hsin-Ping Chiu, Yao Yu Yeo, Tsoi Ying Lai, Chuan-Tien Hung, Shreyas Kowdle, Griffin D Haas, Sizun Jiang, Weina Sun, Benhur Lee

## Abstract

Coronavirus (CoV) Nsp15 is a viral endoribonuclease (EndoU) with a preference for uridine residues. CoV Nsp15 is an innate immune antagonist which prevents dsRNA sensor recognition and stress granule formation by targeting viral and host RNAs. SARS-CoV-2 restricts and delays the host antiviral innate immune responses through multiple viral proteins, but the role of SARS-CoV-2 Nsp15 in innate immune evasion is not completely understood. Here, we generate an EndoU activity knockout rSARS-CoV-2^Nsp15-H234A^ to elucidate the biological functions of Nsp15. Relative to wild-type rSARS-CoV-2, replication of rSARS-CoV-2^Nsp15-H234A^ was significantly decreased in IFN-responsive A549-ACE2 cells but not in its STAT1 knockout counterpart. Transcriptomic analysis revealed upregulation of innate immune response genes in cells infected with rSARS-CoV-2^Nsp15-H234A^ relative to wild-type virus, including cGAS-STING, cytosolic DNA sensors activated by both DNA and RNA viruses. Treatment with STING inhibitors H-151 and SN-011 rescued the attenuated phenotype of rSARS-CoV-2^Nsp15-H234A^. SARS-CoV-2 Nsp15 inhibited cGAS-STING-mediated IFN-β promoter and NF-κB reporter activity, as well as facilitated the replication of EV-D68 and NDV by diminishing cGAS and STING expression and downstream innate immune responses. Notably, the decline in cGAS and STING was also apparent during SARS-CoV-2 infection. The EndoU activity was essential for SARS-CoV-2 Nsp15-mediated cGAS and STING downregulation, but not all HCoV Nsp15 share the consistent substrate selectivity. In the hamster model, rSARS-CoV-2^Nsp15-H234A^ replicated to lower titers in the nasal turbinates and lungs and induced higher innate immune responses. Collectively, our findings exhibit that SARS-CoV-2 Nsp15 serves as a host innate immune antagonist by targeting host cGAS and STING.

**Significance statement:** Host innate immune system serves as the primary defense against pathogens, including SARS-CoV-2. Co-evolving with the hosts, viruses develop multiple approaches to escape the host surveillance. SARS-CoV-2 silences and dysregulates innate immune responses, and the chaos of antiviral IFN responses highly correlates to COVID-19 disease severity. Nsp15 is a conventional innate immune antagonist across coronaviruses, but the biological impact about SARS-CoV-2 Nsp15 is still unclear. Here, we provide a novel insight that SARS-CoV-2 Nsp15 hampers the expression of innate immune regulator – cGAS and STING via its endoribonuclease activity, then further ameliorates virus replication.

## Introduction

Severe acute respiratory syndrome coronavirus 2 (SARS-CoV-2) is a betacoronavirus that is the cause of coronavirus disease 2019 (COVID-19). Previous coronaviruses infecting humans, including HCoV-229E, OC43, HKU1, NL63, as well as the highly pathogenic SARS-CoV and MERS-CoV, have resulted in respiratory illnesses ranging from mild to lethal, depending on viral genetic diversity and host-specific factors.

Coronaviruses (CoVs) are enveloped, positive-sense, and single-stranded RNA viruses with genome approximately 30 kilobases in length, featuring 5′-capping and 3′-polyadenylation. The viral genome RNA encodes multiple open reading frames (ORFs), responsible for translation of nonstructural replicase proteins (Nsp1-16), structural proteins (spike, membrane, envelope, nucleocapsid), and several accessory proteins (1). Nsp15 is a uridine-specific endoribonuclease with preference for cleaving RNA substrates 3′ of uridines. *In vitro* cleavage assays demonstrate that SARS-CoV-2 Nsp15 selectively targets the unpaired uridine within structurally unstable RNA and has preference for purines 3′ of the cleaved uridine (2). Mouse hepatitis virus (MHV) Nsp15 not only cleaves positive sense genomic RNA with strong preference for U↓A and C↓A sequences, but also targets the 5′ poly(U) tract in negative sense viral RNA during MHV infection (3, 4). SARS-CoV-2 Nsp15 targets dsRNA and preferentially degrades AU-rich dsRNA through its dsRNA nickase activity (5). In addition to viral targets, porcine epidemic diarrhea virus (PEDV) Nsp15 degrades porcine TBK1 and IRF3 dependent on its EndoU activity (6).

The innate immune system plays pivotal roles on sensing virus infection and evoking initial antiviral responses. The interferon (IFN)-associated responses and the expression of interferon-stimulated genes (ISGs) constitute the major front-line of defense. SARS-CoV-2 infection delays and limits IFNs and ISGs responses especially at early stage of viral replication, and this dysregulation of antiviral innate immune responses contributes to the severity of COVID-19 (7, 8). SARS-CoV-2 has evolved different strategies to interfere with innate immune responses or otherwise co-opt the host cell’s machinery to facilitate optimal viral replication (9). Nsp15 is a conserved host innate immune antagonist across coronaviruses. Nsp15 tampers the recognition of viral RNA by cytosolic dsRNA sensors such as MDA5, PKR, and OAS, that are integral to the antiviral defense (10, 11). Furthermore, Nsp15 also prevents stress granule assembly and cell apoptosis in macrophages by controlling the accumulation of viral dsRNA intermediates and shortening the poly(U) sequences in viral RNA (3, 11, 12); well-defined pathogen-associated molecular patterns (PAMPs) sensed by the host pattern recognition receptors (PRRs). Nsp15 is considered a virulence determinant as CoVs such as MHV, PEDV, and avian infectious bronchitis virus (IBV) engineered to express a catalytically inactive Nsp15 mutant exhibit an attenuated phenotype *in vitro* and *in vivo* (11, 13, 14).

The vast majority of studies on SARS-CoV-2 Nsp15’s ability to antagonize the production of type I IFNs and downstream signaling have relied solely on *in vitro* biochemical and IFN-β promoter and IFN-stimulated response element (ISRE) reporter assays (15, 16). Recently, Weiss and colleagues showed that a recombinant SARS-CoV-2 with a catalytically inactive Nsp15 mutant had impaired replication kinetics in primary human nasal epithelial cells due to increased activation of IFN responses and PKR pathway (17). Nevertheless, the molecular mechanism linked to Nsp15 EndoU activity remains underexplored. To better understand the biological significances of SARS-CoV-2 Nsp15 in the context of viral infection, we generated recombinant SARS-CoV-2 viruses with deficient or absent EndoU activity. Here, we elucidated the replication phenotypes and transcriptomic signatures during wild-type and Nsp15 mutant virus infection and found cGAS and STING as host targets of Nsp15. We further physiologically evaluated the pathogenesis of Nsp15 EndoU inactive SARS-CoV-2 in hamsters.

## Results

### Attenuation of Nsp15 catalytically mutant SARS-CoV-2 in IFN-competent human lung-derived epithelial cell lines

To investigate the biological functions of SARS-CoV-2 Nsp15 during viral replication, we employed the bacterial artificial chromosome (BAC) system to generate recombinant SARS-CoV-2 (rSARS-CoV-2) expressing Venus reporter with two distinct Nsp15 mutations: H234A and N277A (Fig. 1A). The H234A mutation results in a catalytically inactive Nsp15, whereas N277A exhibits lower *in vitro* EndoU activity and specificity for uridine (18, 19). rSARS-CoV-2 bearing wild-type (WT) and mutant (H234A, N277A) Nsp15 replicated equivalently in IFN-deficient Vero E6 cells (Fig. 1B). Immunoprecipitation of infected cell lysates showed that the H234A and N277A mutants were expressed comparably to WT Nsp15 protein during viral infection. (Fig. 1C). To examine the effects of IFN signaling and downstream ISGs on the replication of WT versus mutant Nsp15 viruses, we used A549-ACE2 cells and its isogenic A549-ACE2/STAT1 KO counterpart that is deficient in IFN signaling (20) (Fig. 1D). Relative to WT virus, the replication of the Nsp15_H234A_ mutant virus was markedly attenuated in IFN-competent A549-ACE2 cells (Fig. 1E). However, in A549-ACE2/STAT1 KO cells, Nsp15_H234A_ mutant virus achieved peak titers comparable to WT virus despite a lag at earlier time points (Fig. 1F). Importantly, the replication of each virus was elevated in STAT1 KO cells, confirming the sensitivity of SARS-CoV-2 to IFN responses, which have been described previously (21–23). Area under the curve (AUC) analysis of the viral growth trajectories quantifies the significantly attenuated phenotype of the Nsp15_H234A_ mutant virus in A549-ACE2 cells, and the enhanced replication of each virus in STAT1 KO cells with the greatest enhancement seen for the Nsp15_H234A_ mutant virus (Fig. 1G). Collectively, these results suggest that Nsp15 from SARS-CoV-2 functions as a negative regulator of IFN-mediated antiviral responses.

**Fig. 1.**
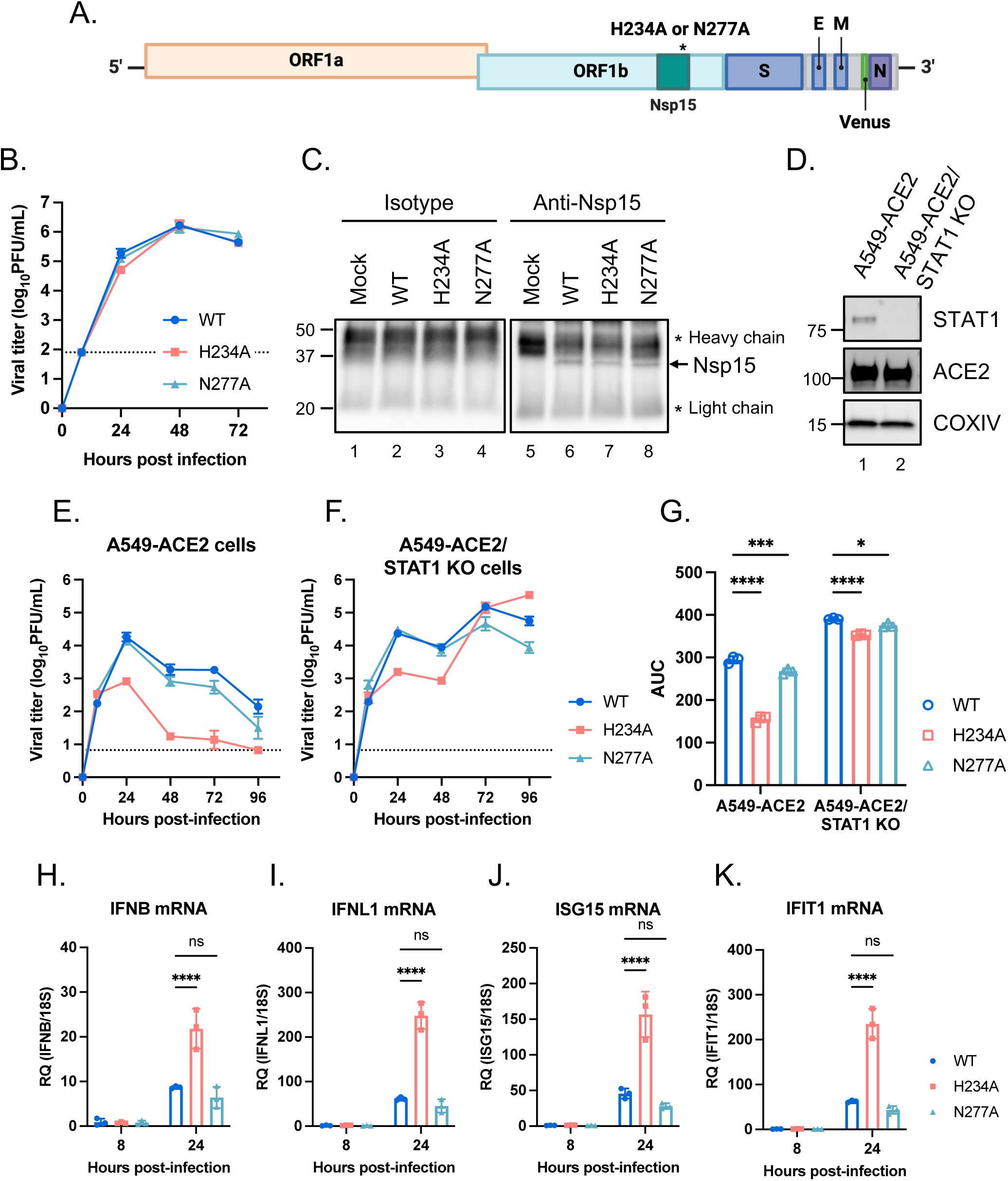
Infection of Nsp15 wild-type and mutant SARS-CoV-2 in Vero E6 and A549-ACE2 cells. (A) Schematic diagram of the recombinant SARS-CoV-2 genome with Nsp15 H234A and N277A mutations. (B) Replication kinetics of Nsp15_WT_, Nsp15_H234A_, and Nsp15_N277A_ rSARS-CoV-2 in Vero E6 cells (MOI of 0.001). (C) Cell lysates from mock- and virus-infected Vero E6 cells (MOI of 0.001) after 48 h of infection were incubated with anti-Nsp15 antibody or isotype control. Immunoblot analysis applied for the immunoprecipitated Nsp15 proteins. (D) Western blot for ACE2 and STAT1 in A549-ACE2 cells with or without STAT1 knockout. (E and F) Replication kinetics of Nsp15_WT_, Nsp15_H234A_, and Nsp15_N277A_ rSARS-CoV-2 in A549-ACE2 and A549-ACE2/STAT1 KO cells (MOI of 1). (G) The area under the curve (AUC) was measured from each viral growth curve and plotted as a bar graph. Data are mean ± SD (n = 3) and analyzed by two-way ANOVA with Dunnett’s multiple comparison test. (H-K) A549-ACE2 cells were uninfected or infected by Nsp15_WT_, Nsp15_H234A_, and Nsp15_N277A_ rSARS-CoV-2 at MOI of 1 for 8 and 24 h. Total RNA collected for evaluating the host responses by RT-qPCR. Relative gene expression was normalized by 18S rRNA and presented relative to mock infection. Data are mean ± SD (n = 3) and analyzed by two-way ANOVA with Dunnett’s multiple comparison test. * *P*≤0.05; *** *P*≤0.001; **** *P*≤0.0001; ns, not significant.

### Enhancement of innate immune responses during Nsp15_H234A_ SARS-CoV-2 infection

Nsp15 is well-documented to enable the escape of cytosolic dsRNA sensors recognition, including PKR and OAS (24). The results shown in Fig. 1E and 1F also implicated the importance of IFN-induced signaling pathways in controlling Nsp15_H234A_ mutant virus infection. Distinct from Nsp15_WT_ and Nsp15_N277A_ viruses, which exhibited similar replication dynamics, Nsp15_H234A_ virus with attenuated phenotype increased IFNs and ISGs expression in A549-ACE2 cells (Fig. 1H-1K), implying that the replication damage observed during Nsp15_H234A_ virus infection is a consequence of enhanced innate immune responses. Additionally, we monitored the PKR and eIF2α phosphorylation, as well as rRNA degradation, which serve as indicators of PKR and OAS/RNase L activation described previously. At 8 hours post-infection (hpi), A549-ACE2 cells infected with Nsp15_H234A_ virus exhibited increased levels of pPKR, peIF2α, and decay of 28S and 18S rRNA compared to those of the other two viruses. However, by 24 hpi, these elevations were reversed, probably due to significant decrease in replication shown in the Nsp15_H234A_ mutant (Fig. S1).

### Beneficial role of Nsp15 EndoU activity in VSV replication

To determine if the innate immune evasion properties of Nsp15 apply in the context of non-CoV infections, we introduced FLAG-tagged WT and H234A Nsp15 into recombinant vesicular stomatitis virus (rVSV) expressing EGFP reporter (Fig. 2A). The expression of Nsp15-FLAG was verified in Vero cells infected with rVSV-Nsp15_WT_ and -Nsp15_H234A_ (Fig. 2B). We next infected A549-ACE2 and A549-ACE2/STAT1 KO cells with parental rVSV-EGFP and rVSV bearing WT and H234A Nsp15. Regardless of STAT1 knockout status, parental rVSV-EGFP outperformed those expressing either WT or H234A Nsp15, thus indicating the insertion of SARS-CoV-2 Nsp15 in the 3′ end of the viral genome is not a gain-of-function modification for VSV. Nonetheless, rVSV-Nsp15_WT_ had a small but significant growth advantage over rVSV-Nsp15_H234A_ in A549-ACE2 cells but not in A549-ACE2/STAT1 KO cells (Fig. 2C and 2D). Furthermore, rVSV-Nsp15_H234A_ infection triggered higher expression of ISGs and inflammatory cytokines than rVSV-Nsp15_WT_ infection (Fig. 2E-2H), corroborating the features of Nsp15 about innate immune antagonism.

**Fig. 2.**
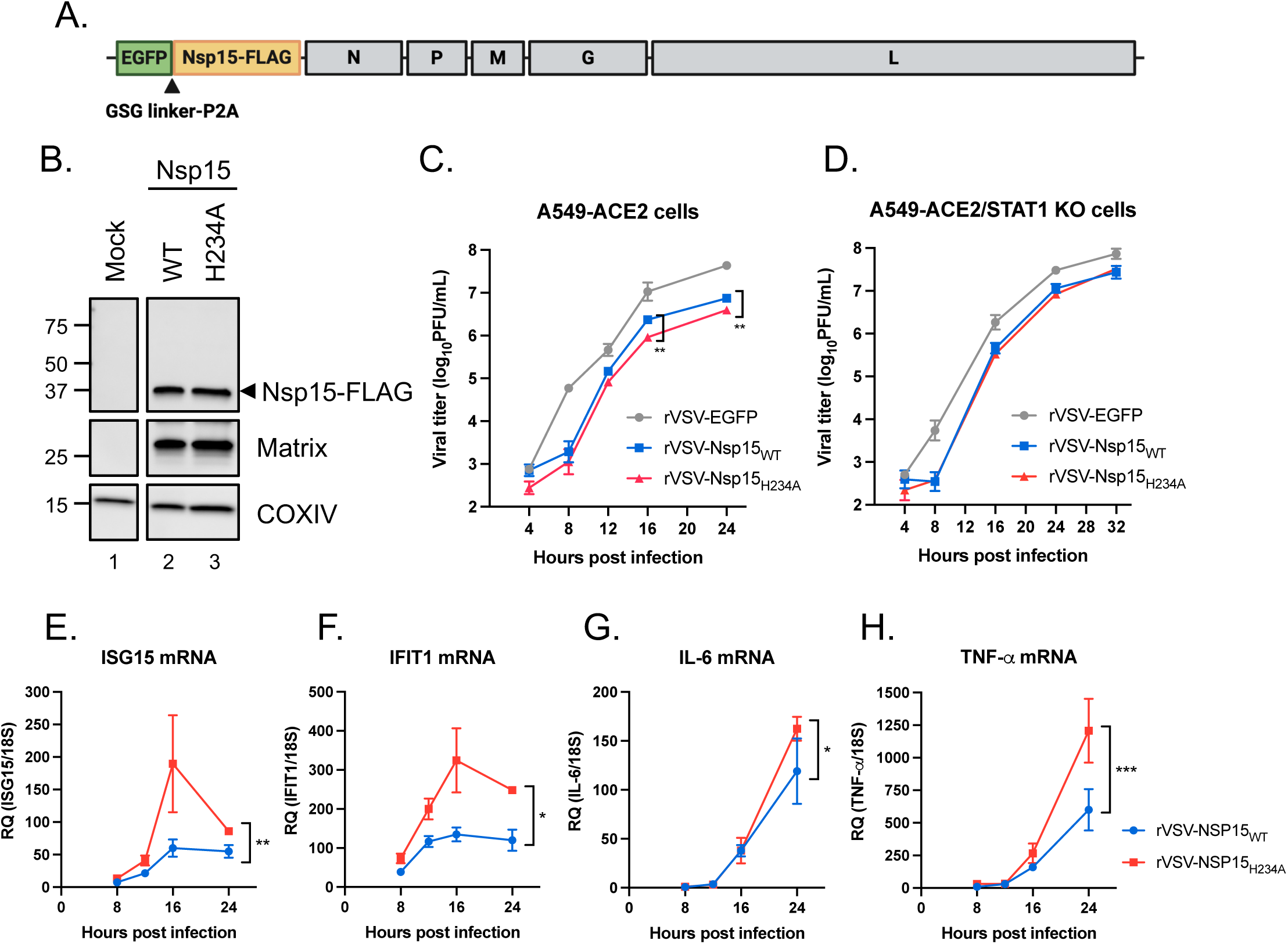
Innate immune antagonism by Nsp15 during VSV infection. (A) Design of rVSV-EGFP genome bearing SARS-CoV-2 Nsp15. (B) Vero cells were uninfected or infected by rVSV-EGFP expressing WT and H234A Nsp15 at MOI of 0.1 for 16 h. Western blot performed to verify the expression of indicated proteins. (C and D) Replication kinetics of parental rVSV-EGFP and rVSV-EGFP expressing WT and H234A Nsp15 in A549-ACE2 and A549-ACE2/STAT1 KO cells (MOI of 0.1). Data are mean ± SD (n = 3) and analyzed by two-way ANOVA with Dunnett’s multiple comparison test. (E-H) Total RNA collected from A549-ACE2 cells with rVSV-Nsp15_WT_ and -Nsp15_H234A_ infection and mock infection. RNA level of indicated host genes relative to 18S rRNA was measured by RT-qPCR and presented by fold change over mock infection. Data are mean ± SD (n = 3) and analyzed by two-way ANOVA. * *P*≤0.05; ** *P*≤0.01; *** *P*≤0.001.

### Nsp15 antagonizes host antiviral innate immune responses and dampens cellular metabolism

To explore the host cellular responses modulated by Nsp15 during viral infection, we infected A549-ACE2 cells with rSARS-CoV-2 harboring WT, H234A, or N277A Nsp15, as well as a mock infection control. Cells were collected at 8 or 24 hpi for poly(A) enriched bulk RNA sequencing (RNA-seq) (Fig. 3A). Principal component analysis (PCA) revealed that cells infected with the Nsp15_H234A_ mutant virus exhibited a unique transcriptional profile, distinct from both mock-infected cells and cells infected with the other two viruses. In contrast, the transcriptional profiles from Nsp15_WT_ and Nsp15_N277A_ virus infection were closely clustered, implying similar transcriptional gene programs (Fig. 3B). There were more unique differentially expressed genes (DEGs) in Nsp15_H234A_ virus-infected cells than Nsp15_N277A_ virus-infected cells when compared to Nsp15_WT_ virus infection, and the contrast increased from 8 to 24 hpi (Fig. 3C), suggesting the key role of Nsp15 catalytic activity toward divergent gene expression programs in SARS-CoV-2-infected cells. We next performed gene set expression analysis (GSEA) between Nsp15_WT_ and Nsp15_H234A_ or Nsp15_N277A_ virus-infected cells using the Molecular Signature Database (MSigDB) Hallmark gene sets (25–27). At 8 hpi, transcripts from Nsp15_H234A_ virus-infected cells were significantly enriched in innate immune (IFN signaling) and metabolic (oxidative phosphorylation and MYC targets) signatures, but no observable changes were found in Nsp15_N277A_ virus-infected cells (Fig. 3D, left). At 24 hpi, transcripts from Nsp15_H234A_ viral infection were significantly higher in other metabolic-associated signatures (e.g. mTORC1 signaling and glycolysis), while transcriptional signature from Nsp15_N277A_ virus-infected cells began to resemble the Nsp15_H234A_ viral infection at 8 hpi (Fig. 3D, right). We further performed gene set variation analysis (GSVA) using antiviral innate immune and cellular metabolism molecular signatures from the MSigDB Gene Ontology gene sets (28). Antiviral innate immune signatures were consistently higher in Nsp15_H234A_-infected cells at 8 hpi and further elevated at 24 hpi; these pathways were generally lower in Nsp15_N277A_ virus infection and lowest in Nsp15_WT_ virus infection across both time points. Cellular respiration signatures, while highest in mock-infected cells, were less dampened in Nsp15_H234A_ virus-infected cells as compared to Nsp15_WT_ and Nsp15_N277A_ virus-infected cells, and the contrasts are much larger at 24 hpi (Fig. 3E). Our findings suggest that the catalytic inactivity of Nsp15 leads to the promotion of antiviral innate immune responses and retention of cellular metabolism. Consistent with GSVA data, Nsp15_H234A_ virus infection showed noticeably higher expression of genes associated with key components of cellular respiration and antiviral innate immune responses compared to Nsp15_WT_ or Nsp15_N277A_ virus infection (Fig. 3F). Of key interest is *CGAS* and *STING1* that are robustly expressed by 24 hpi, as their gene products serve as central regulators of DNA-mediated innate immune responses, but they have also been implicated in innate immune responses during RNA virus infection, including SARS-CoV-2 (29, 30).

**Fig. 3.**
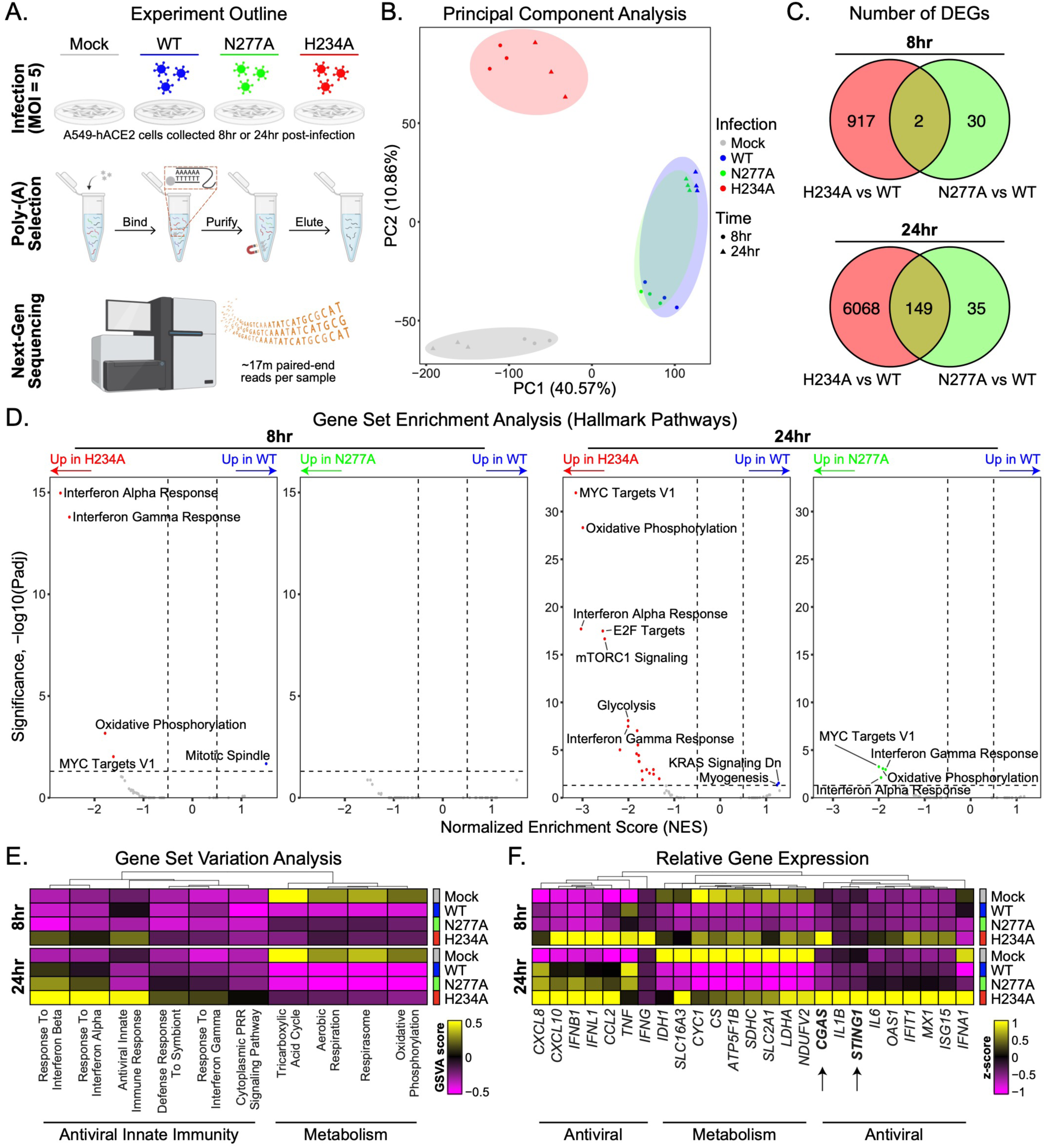
The global transcriptional signatures from Nsp15 wild-type and mutant SARS-CoV-2 infected A549-ACE2 cells. A549-ACE2 cells with mock, Nsp15_WT_, Nsp15_H234A_, and Nsp15_N277A_ rSARS-CoV-2 infection for 8 and 24 h (MOI of 5). Total RNA with poly(A) enrichment followed by RNA sequencing analysis. (A) Schematic of bulk RNA-seq experimental design (n = 3 per group). (B) PCA of total normalized transcript abundance from mock, Nsp15 wild-type and mutant rSARS-CoV-2 infection. Sparse PCA depicts the global transcriptome of individual sample. (C) Venn diagram for unique and shared differentially expressed genes (Padj < 0.05 & |log2FC| > 1) in cells infected with Nsp15_H234A_ and Nsp15_N277A_ mutants compared to Nsp15_WT_ virus. (D) Volcano plots showing GSEA results generated using MSigDB Hallmark pathways. (E) Cluster heatmap of GSVA scores generated using representative innate immune and metabolic signatures from MSigDB Gene Ontology signatures. (F) Expression heatmap of representative innate immune and metabolic genes in across mock, wild-type, and mutant rSARS-CoV-2 infection.

### Decline of cGAS and STING during SARS-CoV-2 infection

Given that cGAS and STING are potential host targets of Nsp15 from the RNA-seq results (Fig. 3F), we orthogonally verified their expression during viral infection. At 24 hpi, mRNA levels of cGAS and STING were decreased in A549-ACE2 cells infected with Nsp15_WT_ rSARS-CoV-2 relative to those infected with Nsp15_H234A_ rSARS-CoV-2 (Fig. 4A and 4B). To assess the protein levels, we conducted infection of Nsp15_WT_ and Nsp15_H234A_ viruses in hamster BHK-21-ACE2 cells with human cGAS and STING overexpression. BHK-21 cells are preferred due to their transfection efficiency and innate immune deficiency (31–34), which minimizes growth discrepancies between Nsp15_WT_ and Nsp15_H234A_ rSARS-CoV-2. In these cells, overexpressed cGAS were reduced following Nsp15_WT_ virus infection at 48 hpi relative to mock-infected cells (58% reduction) and those infected with Nsp15_H234A_ virus (74% reduction), but the decrease in overexpressed STING was more moderate with 56% and 43% reduction, respectively, for the same conditions (Fig. 4C).

**Fig. 4.**
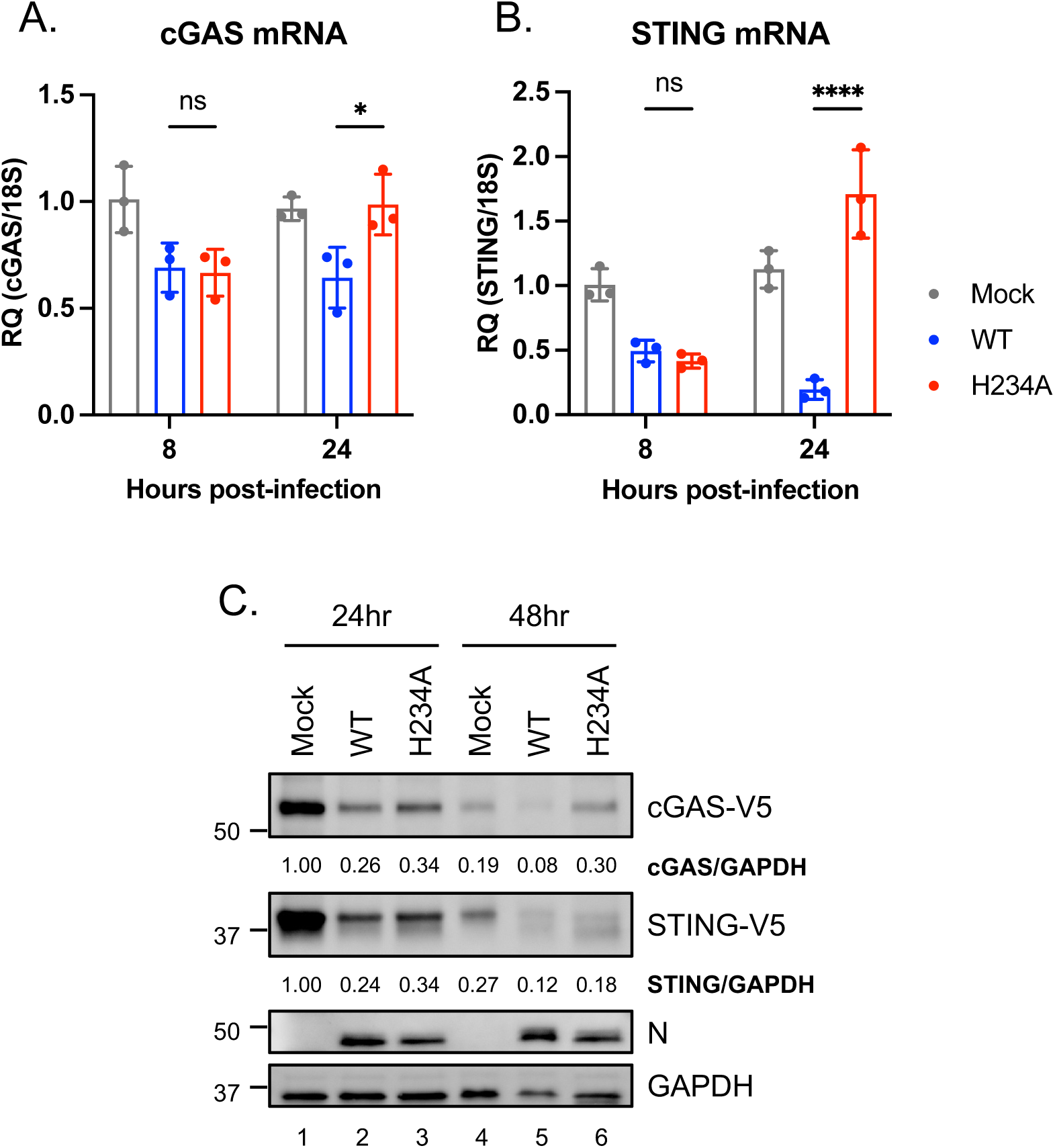
Decrease in cGAS and STING during SARS-CoV-2 infection. (A and B) Endogenous cGAS and STING mRNA levels in A549-ACE2 cells uninfected or infected with Nsp15_WT_ and Nsp15_H234A_ rSARS-CoV-2 at MOI of 5 for 8 or 24 h. RT-qPCR results are presented relative to the expression of 18S rRNA. Data are mean ± SD (n = 3) and analyzed by two-way ANOVA with Tukey’s multiple comparison test. * *P*≤0.05; **** *P*≤0.0001; ns, not significant. (C) BHK-ACE2 cells were mock infected or infected with Nsp15_WT_ and Nsp15_H234A_ rSARS-CoV-2 at MOI of 0.1, further transfected with cGAS and STING plasmids at 4 hpi. At 24 and 48 hpi, cell lysates were harvested for clarifying the indicated protein levels.

### Nsp15 dampens cGAS-STING mediated innate immune responses

To further verify the biological significance of Nsp15 antagonizing the cGAS-STING pathway during virus infection, we infected A549-ACE2 cells with Nsp15_WT_ and Nsp15_H234A_ rSARS-CoV-2 in the presence or absence of STING inhibitors, H-151 or SN-011, chosen for their distinct mechanisms of STING inhibition (35, 36). Prior to viral infection, we preliminarily carried out WST-1 assay to evaluate the cell toxicity of chemicals within the range for further experiments (Fig. 5A and 5B, black dot line). Under STING inhibitor treatment, Nsp15_H234A_ viral replication was enhanced by 10-20-fold, whereas Nsp15_WT_ viral titer remained unaffected (Fig. 5A and 5B, red versus blue lines), emphasizing the role of Nsp15 in antagonizing the cGAS-STING pathway.

**Fig. 5.**
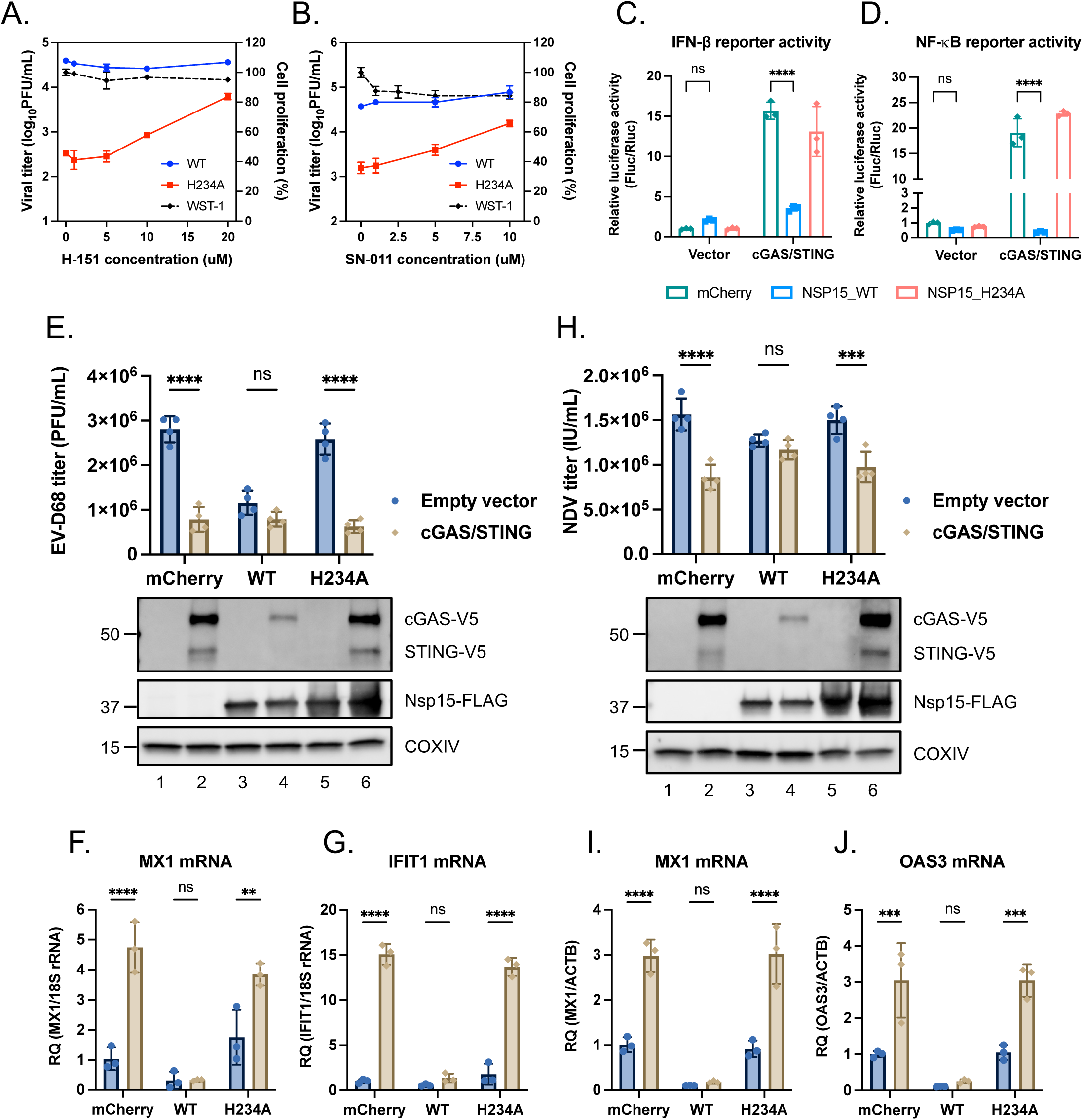
Inhibition of host cGAS-STING pathway by Nsp15. (A and B) A549-ACE2 cells were pretreated with H-151 or SN-011 for 16 h prior to viral infection. Cells were then infected with Nsp15_WT_ (blue line) and Nsp15_H234A_ (red line) rSARS-CoV-2 at MOI of 1 under H-151 or SN-011 treatment for 48 h. Supernatants harvested for viral titration by plaque assay. Cell viability determined by WST-1 assay after incubating with H-151 or SN-011 for 72 h (black dot line). Data are mean ± SD (n = 3). (C and D) HEK293T cells were co-transfected with IFN-β promoter reporter plasmid (C) or NF-κB responsive element reporter plasmid (D), Renilla control plasmid plus the indicated plasmids containing empty vector, cGAS/STING, mCherry and WT and H234A Nsp15 for 48 h. Relative luciferase activity was performed by use of Dual-Glo Luciferase System. Data are mean ± SD (n = 3) and analyzed by two-way ANOVA with Dunnett’s multiple comparison test. (E, F, G) HEK293T transfected with different combinations of plasmids were infected with EV-D68 at MOI of 0.1 for 48 h. Culture supernatants applied for viral titer quantification by plaque assay (E, upper bar graph). Cell lysates collected for protein level determination (E, lower blot graph) and total RNA collected for RT-qPCR (F, G). (H, I, J) HEK293T transfected with different combinations of plasmids were infected with NDV-EGFP at MOI of 1 for 48 h. Culture supernatants applied for viral titer quantification by focus forming assay targeting EGFP (H, upper bar graph). Cell lysates collected for protein level determination (H, lower blot graph) and total RNA collected for RT-qPCR (I, J). Relative target RNA level normalized with that of 18S rRNA or ACTB was shown. Data are mean ± SD (n = 3 or 4). and analyzed by two-way ANOVA with Šídák multiple comparison test. ** *P*≤0.01; *** *P*≤0.001; **** *P*≤0.0001; ns, not significant.

cGAS-STING is a cytosolic DNA-sensing pathway to drive innate immune responses (37), which has been reported to restrict RNA virus infection probably due to mitochondrial DNA leakage during viral infection (38–40). To ascertain the potential that Nsp15 inhibits cGAS-STING-induced innate immune responses, we performed IFN-β promoter and NF-κB reporter assays. In the presence of vector control, Nsp15_WT_ did not alter reporter activity; with cGAS-STING stimulation, it greatly restricted IFN-β promoter and NF-κB reporter activity compared with mCherry control and Nsp15_H234A_ (Fig. 5C and 5D). Moreover, we examined the ability of Nsp15 to facilitate other RNA virus replication by diminishing the antiviral effects induced by cGAS-STING pathway. To this end, HEK293T cells were transfected with cGAS and STING plasmids along with either WT or H234A Nsp15 plasmids, followed by enterovirus D68 (EV-D68) or Newcastle disease virus (NDV) infection. In contrast to mCherry and H234A Nsp15, WT Nsp15 rescued EV-D68 and NDV viral titer under cGAS-STING stimulation (Fig. 5E and 5H). Strikingly, WT Nsp15 reduced EV-D68 viral titer even without cGAS-STING overexpression, but under the same scenario, it had a minimal effect on NDV, probably due to NDV’s encapsidated genome which provided protection from Nsp15. Conversely, under cGAS-STING stimulation, cells with mCherry and H234A Nsp15 expression showing lower viral titer induced more innate immune responses driven by cGAS-STING (Fig. 5F, 5G, 5I, and 5J). Note that cells overexpressing WT Nsp15 exhibited relatively lower levels of cGAS-V5 and STING-V5, revealing that Nsp15 antagonizes cGAS-STING-mediated antiviral responses by downregulating both cGAS and STING.

### Nsp15 EndoU activity is required for cGAS and STING downregulation

The catalytic triad of SARS-CoV-2 Nsp15 (H234, H249, K289) and residues involved in uridine specificity (N277, S293, Y342) have been delineated and characterized through structural and biochemical analyses (18, 19) (Fig. S2). To comprehensively probe the role of these residues in Nsp15’s ability to degrade cGAS and STING mRNAs, we co-transfected WT and the cognate Nsp15 mutant plasmids along with cGAS or STING plasmid into HEK293T cells and assessed RNA and protein levels of cGAS and STING. The mutations located in RNase catalytic triad (H234A, H249A, K289A) abrogated the degradation of cGAS and STING, consistent with the published structural and functional analyses of SARS-CoV Nsp15 (41, 42). N277A and S293A, the mutations involved in uridine discrimination, retained Nsp15’s EndoU activity targeting cGAS and STING (Fig. 6A-6D). Interestingly, S293A was considered a loss-of-function mutation based on an *in vitro* RNA cleavage assay (19); in our hands, the impact of S293A is comparable to that of N277A, which was previously characterized to have lower EndoU activity and uridine specificity *in vitro* (19) but do not appear to have as dramatic effects as the catalytically inactive mutations.

**Fig. 6.**
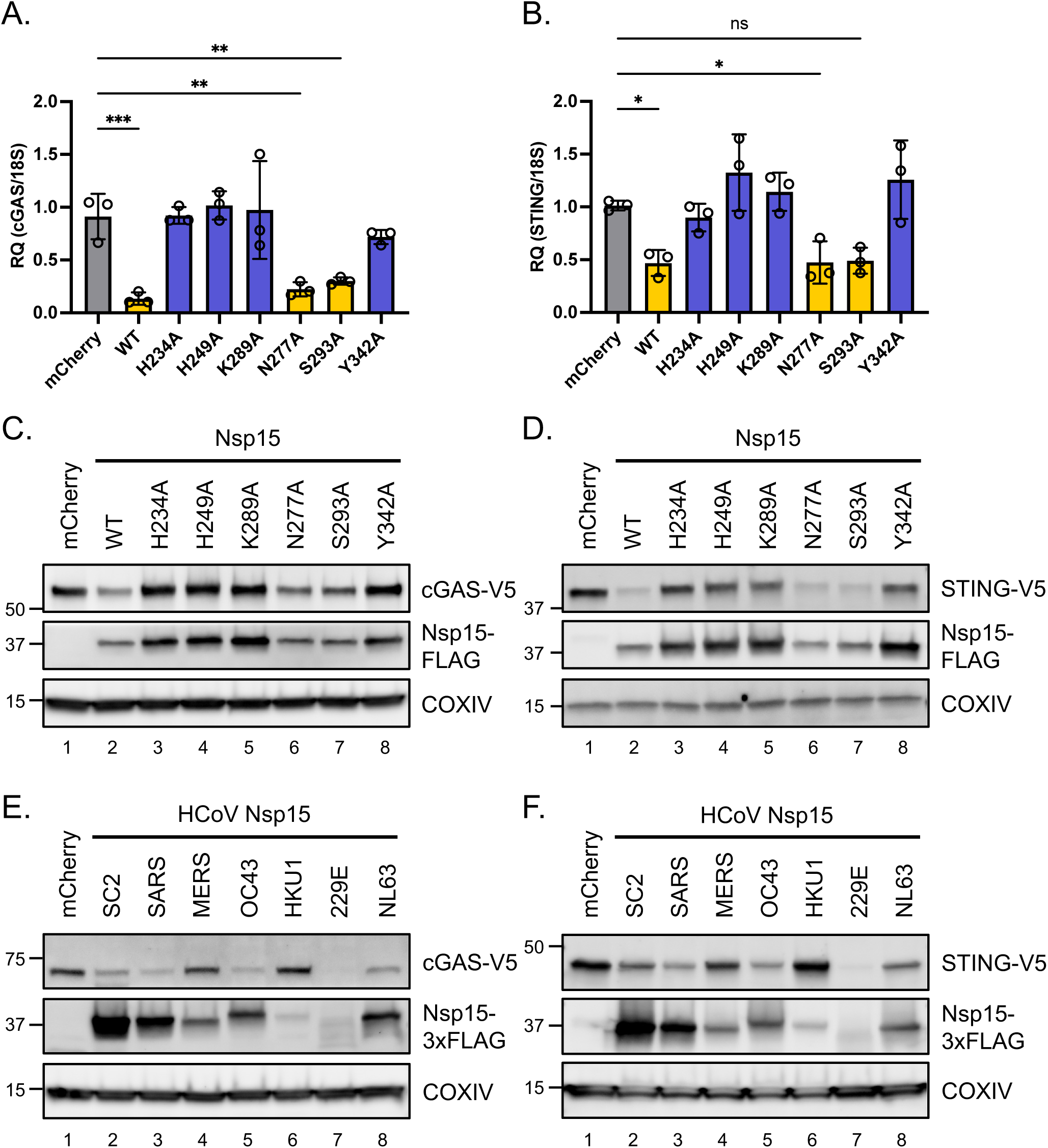
Role of Nsp15 EndoU activity in cGAS and STING downregulation. (A-D) HEK293T cells were transfected with cGAS or STING plasmid along with wild-type and mutant SARS-CoV-2 Nsp15 plasmids for 48 h. RNA levels (A and B) and protein levels (C and D) of cGAS and STING were measured by RT-qPCR and western blot, respectively. RT-qPCR results are presented relative to the expression of 18S rRNA. Data are mean ± SD (n = 3) and analyzed by one-way ANOVA with Dunnett’s multiple comparison test. (E and F) HEK293T cells were transfected with cGAS or STING plasmid plus mCherry or HCoV Nsp15 plasmids for 48 h. Western blot performed to assess the expression of indicated proteins.

Currently identified human CoVs, belonging to either alpha- or betacoronavirus lineages, have variable Nsp15 sequence identity (Fig. S3). To extend the inspection for Nsp15 from other human CoVs targeting cGAS and STING, we generated the Nsp15 constructs for SARS, MERS, OC43, HKU1, 229E, and NL63. The FLAG-tagged Nsp15 from the various human CoVs exhibited variable levels of expression that did not correlate with their ability to degrade cGAS and STING (Fig. 6E and 6F). For example, Nsp15 from 229E had the lowest expression but also appeared to be the most potent degrader of cGAS and STING. Conversely, Nsp15 from SARS-CoV-2 (SC2 in figure) was expressed much better than that of SARS and OC43, but cGAS and STING was not degraded more. While Nsp15 overexpression can execute non-specific cleavage (43), it is unclear how non-specific cleavage can lead to the patterns of cGAS and STING degradation observed where the most highly expressed Nsp15 from SARS-CoV-2 degraded cGAS and STING less potently than the lowest expressed Nsp15 from 229E (compare lane 2 and 7 in Fig. 6E and 6F). Thus, we posit that our data suggest Nsp15 from different CoVs may have different substrate selectivities or enzymatic activities.

### Attenuation of Nsp15 EndoU inactive SARS-CoV-2 in nasal turbinates and lungs of hamsters

The virulence attenuation and pathogenicity of Nsp15 EndoU mutant MHV, PEDV, and IBV have previously been reported *in vitro* and *in vivo* (11, 13, 14). Nsp15_H234A_ mutant rSARS-CoV-2 was attenuated in cells with functional IFN-associated responses; therefore, we sought to investigate the pathogenicity of Nsp15_H234A_ virus in an animal model. We infected golden Syrian hamsters via the intranasal route with Nsp15_WT_ and Nsp15_H234A_ rSARS-CoV-2, and subsequently monitored the body weight change, viral load in tissues, and host responses, respectively. Up to 5 days post-infection (dpi), hamsters infected with Nsp15_WT_ and Nsp15_H234A_ viruses showed comparable weight loss (Fig. 7A). Despite the unanticipated weight loss in hamsters infected with the Nsp15_H234A_ mutant virus, Nsp15_H234A_ SARS-CoV-2 showed reduced viral titer in respiratory tissues, such as nasal turbinates and lungs (Fig. 7B-7E), and provoked robust innate immune responses in lungs (Fig. 7F-7J), which are consistent with the phenotypes observed *in vitro* (Fig. 1E-1K).

**Fig. 7.**
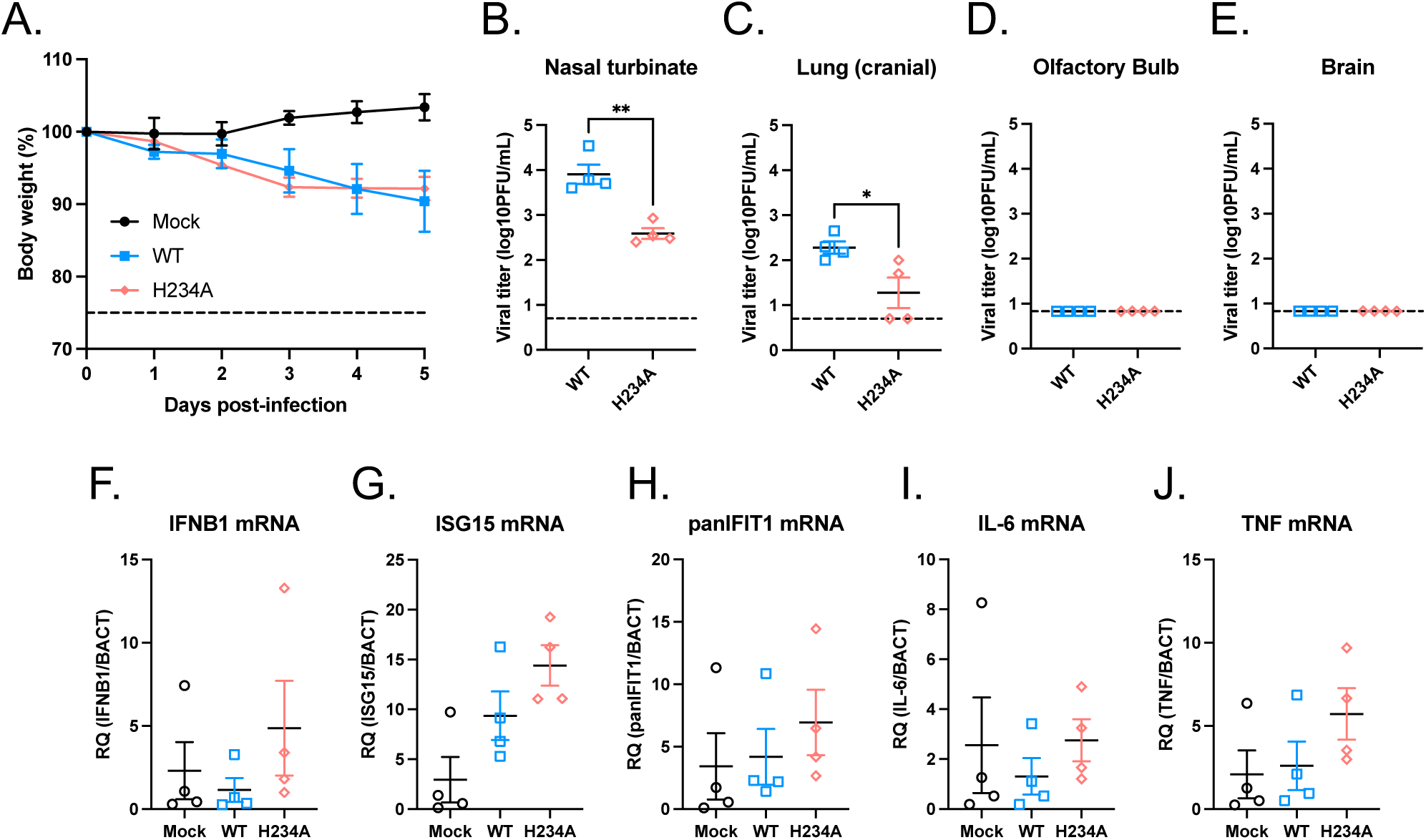
Viral pathogenesis of Nsp15 EndoU inactive SARS-CoV-2 in hamsters. Golden Syrian hamsters were intranasally inoculated with 1x10^5^ PFU of Nsp15_WT_ or Nsp15_H234A_ rSARS-CoV-2, or equivalent volume of PBS. (A) Viral pathogenicity was evaluated by body weight loss. (B-E) Viral titer in the nasal turbinates, lungs, olfactory bulbs, and brains were determined at 5 dpi by plaque assay. Data are mean ± SEM (n = 4) and analyzed by unpaired t-test. * *P*≤0.05; ** *P*≤0.01. (F-J) Host responses in lungs assessed by RT-qPCR. Data are mean ± SEM (n = 4).

## Discussion

Virus infections trigger host innate and adaptive immunity to counteract the spread of foreign pathogens. Meanwhile, viruses have evolved diverse strategies to escape or even manipulate these host defenses. CoV Nsp15, while dispensable for viral survival, significantly affects viral replication and virulence *in vitro* and *in vivo*, as demonstrated in studies on animal CoVs like MHV, PEDV, and IBV (11, 13, 14). Here, we explore the role of SARS-CoV-2 Nsp15 by employing recombinant viruses with functional impairments in the Nsp15 protein. Nsp15 deactivation results in viral attenuation by enhancing host innate immune responses (Fig. 1 and S1). We also found that the host cGAS-STING pathway is a bona fide target of Nsp15 and plays a significant role in the innate immune responses against SARS-CoV-2.

Previous studies have shown that SARS-CoV-2 Nsp15 helps to mediate escape from innate immune surveillance, consistent with the known properties of other CoV Nsp15 proteins (24). Recently, Weiss and colleagues demonstrated in primary nasal epithelial cells that rSARS-CoV-2 with an enzymatically dead Nsp15 was more sensitive to IFN-associated attenuation than its WT counterpart (17). Similar to MHV, SARS-CoV-2 with mutant Nsp15 fails to control dsRNA accumulation, thereby enhancing IFN signaling and PKR activation (17, 44). However, the specific molecular mechanism driven by Nsp15 EndoU activity remains elusive. While how Nsp15 EndoU functions to downregulate viral RNA-derived PAMPs is relatively well-understood, it is not known whether Nsp15-mediated cleavage of specific host transcripts also contribute to dampening of the innate immune responses. Here, we provide a novel perspective that SARS-CoV-2 Nsp15 exerts its EndoU activity to strengthen the resistance of SARS-CoV-2 to host innate immunity in part by downregulating cGAS and STING (Fig. 4-6).

As an RNA virus, SARS-CoV-2 primarily activates RNA-sensing mechanisms, such as MDA5, PKR, OAS, and ZAP (45). Additionally, the DNA-mediated cGAS-STING pathway can also be activated by spike protein-induced fusion and mitochondrial DNA leakage during viral infection (30, 46). The induction of cGAS-STING activity is seen not only in cell culture infections, but also in clinical lung samples from COVID-19 patients (29, 30). Interestingly, pharmacological activation of STING by diABZI have shown protective effects against SARS-CoV-2 infection *in vitro* and *in vivo* (47–49), underscoring the importance of cGAS-STING pathway in host defense against SARS-CoV-2. Our study now directly implicates Nsp15’s EndoU activity in ameliorating the antiviral effects of cGAS-STING activation during SARS-CoV-2 infection; two distinct STING inhibitors (H-151 and SN-011) partially restored attenuated Nsp15_H234A_ SARS-CoV-2 replication, enhancing H234A mutant virus titers by 10-20-fold (Fig. 5A and 5B) and SARS-CoV-2 Nsp15 downregulated cGAS-STING mediated IFN responses and NF-κB activation in an EndoU activity dependent manner (Fig. 5C and 5D). Increasing evidence indicates that the cGAS-STING pathway plays an important role in suppressing SARS-CoV-2 replication given the number of viral proteins that specifically target this pathway. Nsp5 (3CL) suppresses the K63-linked ubiquitination of STING and inhibits recruitment of TBK1 and IKKβ (50). ORF3a blocks cGAS-STING induced autophagy by directly disrupting STING-LC3 interaction (51). ORF10 also restrains cGAS-STING mediated IFN responses as well as autophagy by preventing STING trafficking (52).

Mitochondrial functions and cellular metabolic processes are crucial in regulating innate immune responses. The mitochondrial antiviral signaling protein (MAVS), located on the outer membrane of mitochondria, mitochondria-associated ER membranes (MAMs), and peroxisomes, acts as a vital adaptor for RIG-I and MDA5-mediated antiviral signaling (53). Other mitochondrial-localized proteins such as Tom70, MFN1, and MFN2, are also involved in modulating innate immunity (54). Reactive oxygen species (ROS), a byproduct of oxidative phosphorylation in mitochondria, are closely linked to inflammatory responses and cellular/tissue damage. ROS directly activate NLRP3 inflammasome, triggering inflammatory signaling (55, 56). Additionally, ROS have an antiviral effect by promoting IFN-λ production in influenza-infected human nasal epithelial cells (57). Succinate, an intermediate metabolite of tricarboxylic acid (TCA) cycle, can stabilize HIF-1α and drive proinflammatory gene expression in macrophages (58). In Fig. 3D, SARS-CoV-2 infection with viruses bearing Nsp15_WT_ but not Nsp15_H234A_ suppressed mitochondrial metabolic pathways including oxidative phosphorylation (OXPHOS) and TCA cycle. Our findings in terms of mitochondrial metabolism downregulation in Nsp15_WT_ virus-infected cells aligns with reports that SARS-CoV-2 reduces the expression of OXPHOS genes and proteins in patient samples, hamster model, and K-18 ACE2 mouse model (59, 60). Beyond targeting innate immune pathways directly, SARS-CoV-2 Nsp15 may also impair cellular mitochondrial metabolism to strengthen its inhibitory effect on host antiviral innate immunity. How Nsp15’s EndoU activity contributes directly to this metabolic dysregulation phenotype remains to be determined.

Evidence from epidemiological and modeling studies also implicate Nsp15’s EndoU activity as a contributing factor to SARS-CoV-2 fitness. While most mutations associated with fitness have been ascribed to the spike protein, an analysis of 6.4 million SARS-CoV-2 genomes as of Jan 2022 computationally predicted that Nsp15-T112I (virus level numbering), a marker for the Omicron variant, was independently associated with increased fitness (61). Biochemical analysis subsequently showed that the T113I (expressed ORF numbering) mutation gained a 250% increase in EndoU activity compared to its parental Wuhan isolate sequence. Structural analyses suggest that this mutant enhances Nsp15 hexamer formation and facilitates binding of longer RNA substrates (62). Analysis of the GSAID database showed that while Nsp15-T112I comprise only 0.74% of sequences in December 2021, it is now present in almost 98% of sequences in June 2024. This is a higher prevalence than Omicron defining spike mutations such as E484K (90%) and P681R (95%). Conversely, while H234Y, an EndoU-deficient mutant, first appeared in a subclade of Delta (63), it is only detected in 0.18% of all sequences as of June 2024. These epidemiological analyses underscore the biological significance of Nsp15 to SARS-CoV-2 survival.

Overall, our findings highlight the importance of SARS-CoV-2 Nsp15 in facilitating viral replication and counteracting cellular innate immune responses like cGAS-STING. The attenuated replication observed with Nsp15 EndoU-inactive SARS-CoV-2 in hamster respiratory tissues, coupled with the interaction between Nsp15 and host RNA may offer novel approaches for developing live-attenuated vaccines and antivirals against current and future coronavirus infection.

## Materials and Methods

### Cell lines, viruses, and chemicals

HEK 293T (ATCC, CRL-3216), Vero (ATCC, CCL-81), Vero E6 (ATCC, CRL-1586), and BHK-21-hACE2 cells were maintained in Dulbecco’s modified Eagle’s medium (DMEM) supplemented with 10% fetal bovine serum (FBS) (R&D Systems) and 1% penicillin-streptomycin (P/S) (Gibco) at 37 °C with 5% CO_2_. BHK-21-hACE2 cell, a derivative of BHK-21 cell (ATCC, CCL-10), was established by transduction of lentiviral particle bearing human ACE2 (GeneCopoeia, EX-U1285-Lv105) and selected under 10 ug/mL of puromycin (Gibco). A549-hACE2 and A549-hACE2/STAT1 KO cells (20), gifts from Brad R. Rosenberg, MD, PhD, were cultured in DMEM with 10% FBS and 1% P/S. EV-D68, isolate US/MO/14-18947 (ATCC, VR-1823) were amplified and titrated in RD cells (ATCC, CCL-136). Recombinant rNDV-EGFP LaSota strain (GenBank: KY295917) was amplified in embryonic chicken eggs and titrated in Vero cells. Recombinant rVSV-EGFP Indiana strain was amplified and titrated in Vero cells. STING inhibitors H-151 and SN-011 were purchased from Invivogen and MedChemExpress, respectively.

### Plasmids

The codon optimized Nsp15 open reading frames from SARS-CoV-2 (Wuhan-Hu-1 strain, GenBank: NC045512), SARS-CoV (Urbani strain, GenBank: AY278741), MERS-CoV (EMC/2012 strain, GenBank: NC019843), HCoV-229E (ATCC VR-740, GenBank: AF304460), HCoV-OC43 (ATCC VR-759, GenBank: AY585228), HCoV-HKU1 (GenBank: NC006577), HCoV-NL63 (Amsterdam 1 strain, GenBank: NC005831) fused with 1x FLAG or 3x FLAG were inserted into the pcDNA5/FRT/TO vector cut by HindIII (New England Biolabs) and XhoI (New England Biolabs) by In-Fusion cloning (Takara Bio). The alanine substitutions (H234A, H249A, K290A, N277A, S293A, Y342A) were generated by site-directed mutagenesis. pcDNA3.1/HA-hcGAS-V5 and pcDNA3.1/HA-hSTING-V5 are the kindly gifts from Chia-Yi Yu, PhD.

### Rescue of recombinant SARS-CoV-2 virus

We acquired the SARS-CoV-2 BAC with Venus reporter (USA-WA1/2000) from Luis Martinez-Sobrido, PhD (64). For establishing Nsp15 mutant BAC, the subcloning strategy was applied. The DNA sequence between Nsp15 to Spike was PCR amplified and cloned into pCR-Blunt II-TOPO vector (Invitrogen), and the Nsp15 mutations (H234A and N277A) were generated by site-directed mutagenesis. The inserts with modifications and the original BAC were digested by BstBI (New England Biolabs) and BsmHI (New England Biolabs) and ligated using T4 DNA ligase (New England Biolabs). About virus rescue, Vero E6 cells (1x10^6^ cells per 6-well) were transfected with 4 μg of BAC construct (Nsp15_WT_, Nsp15_H234A_, and Nsp15_N277A_) using 10 uL of Lipofectamine 2000 (Invitrogen). Next day, the regular medium containing transfection mixture was replaced with infection medium (DMEM supplement with 2% FBS). After monitoring the infection rate by Venus signal until day 5 or 6, the culture supernatants designated as seed stocks (P0) were harvested and stored at -80°C. The recombinant viruses were propagated and titrated in Vero E6 cells using plaque assay.

### Rescue of recombinant VSV virus harboring WT and H234A Nsp15

The pEMC-VSV-EGFP viral genome plasmid and the VSV-N, VSV-P, and VSV-L accessory protein plasmids required for the virus rescue were mentioned previously (65). To create pEMC-VSV-EGFP-P2A-Nsp15-FLAG construct, open reading frame of EGFP-GSG linker-P2A-Nsp15-FLAG (WT and H234A) was in-frame replaced with the original EGFP sequence. BSR-T7 cells (3x10^5^ cells per 6-well) were transfected with the rescue DNA mixture using Lipofectamine LTX (Invitrogen) as described (65, 66). Culture medium containing transfection mixture was replaced with infection medium (DMEM supplement with 2% FBS) the next day, cells were further maintained for 2 to 3 days until EGFP positive syncytia appeared. The rescue supernatants harvested from BSR-T7 cells served as stocks for the following amplification in Vero cells.

### Reverse transcription and real-time quantitative PCR (RT-qPCR)

Total RNA was extracted by use of Direct-zol RNA Miniprep Kit (Zymo Research). Equivalent RNA was reverse-transcribed by random hexamer or oligo(dT) primers with LunaScript RT Master Mix Kit (New England Biolabs). qPCR was run with primers targeting specific genes (Table S1 and Table S2) and Luna Universal qPCR Master Mix (New England Biolabs) at the Bio-Rad CFX96 Real-Time PCR system (Bio-Rad). The relative RNA levels of specific RNA were normalized with 18S rRNA (for human), ACTB (for human), or BACT (for hamster), and calculated by the comparative threshold cycle (ΔΔCT) method.

### Western blot and antibodies

Cells were lysed by RIPA lysis buffer (Pierce) containing protease inhibitors cocktail (Roche) and phosphatase inhibitors cocktail (Roche). Equivalent amounts of proteins determined by the Protein Assay Dye Reagent (Bio-Rad) were separated by reducing SDS-PAGE and transferred to a polyvinylidene difluoride (PVDF) membrane, 0.22 um (Bio-Rad). To avoid the nonspecific antibody reaction, membranes were blocked with phosphate-buffered saline blocking buffer (LI-COR; 927-700001) and then probed with the primary antibodies targeting specific proteins. For secondary antibodies incubation, the membranes were washed and then treated with anti-mouse or anti-rabbit Alexa Fluor 647-conjugated secondary antibodies (Invitrogen). The fluorescent signals were developed using the ChemiDoc MP imaging system (Bio-Rad). The following primary antibodies were applied: mouse anti-SARS N (clone 1C7), which cross-reacts to SARS-CoV-2 N, was provided by James A. Duty, PhD. Rabbit anti-SARS-CoV-2 Nsp15 (GTX135737) from GeneTex. Rabbit anti-ACE2 (ab108252), rabbit anti-pPKR (ab32036) and mouse anti-GAPDH (ab8245) from Abcam. Rabbit anti-STAT1 (#14994), rabbit anti-peIF2α (#3398), rabbit anti-PKR (#12297), rabbit anti-V5 (#13202), mouse anti-FLAG (#8146), and rabbit anti-COXIV (#4850) from Cell Signaling Technology. Rabbit anti-eIF2α (11170-1-AP), rabbit anti-cGAS (26416-1-AP), and rabbit anti-STING (19851-1-AP) from Proteintech.

### Immunoprecipitation

Cells (4 mg of lysates) lysed by RIPA lysis buffer (Pierce) containing protease inhibitor cocktail (Roche) were immunoprecipitated with pre-incubated mixture of 50 µL of protein A/G magnetic beads (Pierce) and 10 µg of anti-SARS-CoV-2 Nsp15 antibody (16820-1-AP, Proteintech) or rabbit IgG (12–370, EMD Millipore) at 4°C for overnight. The antibody-protein complexes were then washed 3 times by Tris-buffered saline (TBS) containing 0.05% Tween 20 and eluted by 50 µL of reducing SDS-PAGE sample buffer at room temperature. The pull-down Nsp15 proteins were clarified by western blot assay.

### Reporter assay

HEK293T cells were co-transfected with Firefly luciferase reporter plasmids under control of the IFN-β promoter or NF-kB responsive elements, and pRL-TK control plasmid along with the indicated plasmids including empty vector, cGAS/STING, mCherry, and Nsp15 (WT or H234A) for 48 h. Relative luciferase activity (Firefly/Renilla) was performed by use of Dual-Glo Luciferase System (Promega).

### Hamster challenge studies

The rSARS-CoV-2 with Venus reporter (Nsp15_WT_ and Nsp15_H234A_) applied for hamster infection were rescued and propagated in Vero-hTMPRSS2 cells (66). Viral titers were determined in Vero E6 cells using plaque assay. Nsp15 sequences were confirmed by Sanger sequencing. Six to eight week-old female Golden Syrian hamsters (HsdHan:AURA) were purchased from Inotiv. For the challenge, hamsters were anesthetized with a ketamine/xylazine cocktail before administration of 50 µL of total volume split between each nostril. Animals were challenged with 1x10^5^ PFU of each virus. A group of healthy control animals was left untreated. Weight changes of the animals were monitored for 5 days. Animals from each group was euthanized at days 5 post-challenge to harvest nasal turbinates, lung lobes, olfactory bulbs, and brain. The nasal turbinates, olfactory bulbs, brain, and each of the upper right (cranial) and lower right (caudal) lung lobes were homogenized in 1 mL of sterile PBS. Viral titers were determined by plaque assay on Vero E6 cells. The homogenates from upper right lung lobes were mixed with TRIzol LS reagent (Invitrogen) for the evaluation of host responses by RT-qPCR.

### Bulk RNA-seq processing and analysis

Total RNA from mock-infected and virus-infected cells was lysed with TRIzol reagent (Invitrogen), then extracted and on-column DNase I treated using Direct-zol RNA Miniprep kit (Zymo Research). RNA samples were delegated to GENEWIZ, lnc. for polyadenylated RNA enrichment, RNA-seq library preparation, and sequencing process. Sequencing libraries were sequenced on an Illumina HiSeq platform (2x150bp, ∼350M pair-end reads). The reference genome was generated by concatenating the hg38 human and rSARS-CoV-2-Venus (termed SC2-Venus) genomes, which was used for aligning fastq reads using STAR (v2.7.11a). Transcript abundances were then quantified using salmon (v1.10.2) and scaled by transcript length and library size using the R package ‘tximport’ (v1.26.1). Principal component analysis (PCA) was performed using the R package ‘stats’ (v4.2.1). Differential gene expression (DEG) was performed using the R package ‘DESeq2’ (v1.38.3); the false discovery rate for Benjamini-Hochberg p-value adjustment was set to 0.05. Gene set enrichment analysis (GSEA) was performed by first ranking the DEGs (scored using log2 fold change × adjusted p-values), then using the R package ‘fgsea’ (v1.24.0) with the default parameters. Gene set variation analysis (GSVA) was performed using the R package ‘GSVA’ (v1.46.0). The gene signatures used for GSEA (Hallmark) and GSVA (Gene Ontology) were obtained from MSigDB using the R package ‘msigdbr’ (v7.5.1). For data visualization, Fig. 4B and 4D were generated using the R packages ‘ggplot2’ (v3.5.0) and ‘ggrepel’ (v0.9.5), Fig. 4C was generated using the R package ‘ggvenn’ (v0.1.10), and Fig. 4E and 4F were generated using the R packages ‘ComplexHeatmap’ (v2.14.0) and ‘circlize’ (v0.4.16).

### Statistical analysis

One-way and two-way AVONA were used to estimate the statistical significance among multiple groups and conditions. Unpaired t-test was used to estimate the statistical significance between two groups. Representative data are shown as mean ± standard deviation (SD) or mean ± standard error of the mean (SEM) with biological triplicates or quadruplicates. *P*≤0.05 was considered statistically significant. * *P*≤0.05; ** *P*≤0.01; *** *P*≤0.001; **** *P*≤0.0001; ns, not significant. Statistical significance was calculated by use of Prism 10 (GraphPad).

## Data availability

All data are included in the manuscript and supporting information. RNA-seq raw and processed data are available at NCBI Gene Expression Omnibus (GEO): GSE274310.

## Author contributions

H-P.C. and B.L. conceived and designed the study. H-P.C., T.Y.L., C-T.H., S.K., G.D.H., and W.S. performed experiments and collected data. Y.Y.Y. conducted the NGS data processing and analysis. H-P.C. and Y.Y.Y. analyzed the data and wrote the original draft of the manuscript. S.J., W.S., and B.L. reviewed and edited the manuscript.

## Acknowledgements

We thank Drs. Brad Rosenberg, Luis Martinez-Sobrido, and Chia-Yi Yu for sharing materials. We thank members of the Lee lab for experimental assistances and feedback of the project. We thank the Jiang lab for helping with bioinformatics analyses. We thank the Sun lab for helping with hamster related studies. H-P.C. was supported by Postdoctoral Research Abroad Program from Ministry of Science and Technology, Taiwan (110-2917-I-564-020). G.D.H. is supported by the National Science Foundation Graduate Research Fellowship Program (Grant No. 1842169). Any opinions, findings, and conclusions or recommendations expressed in this material are those of the author(s) and do not necessarily reflect the views of the National Science Foundation. S.J. is supported by in part by NIH DP2AI171139, P01AI177687, R01AI149672, a Gilead’s Research Scholars Program in Hematologic Malignancies, a Sanofi iAward, the Bill & Melinda Gates Foundation INV-002704, the Dye Family Foundation, and the Bridge Project, a partnership between the Koch Institute for Integrative Cancer Research at MIT and the Dana-Farber/Harvard Cancer Center. B.L. also acknowledges the Ward-Coleman estate for endowing the Ward-Coleman Chairs at the Icahn School of Medicine at Mount Sinai and diverted support from AI149033.

---

**Fig. S1.**
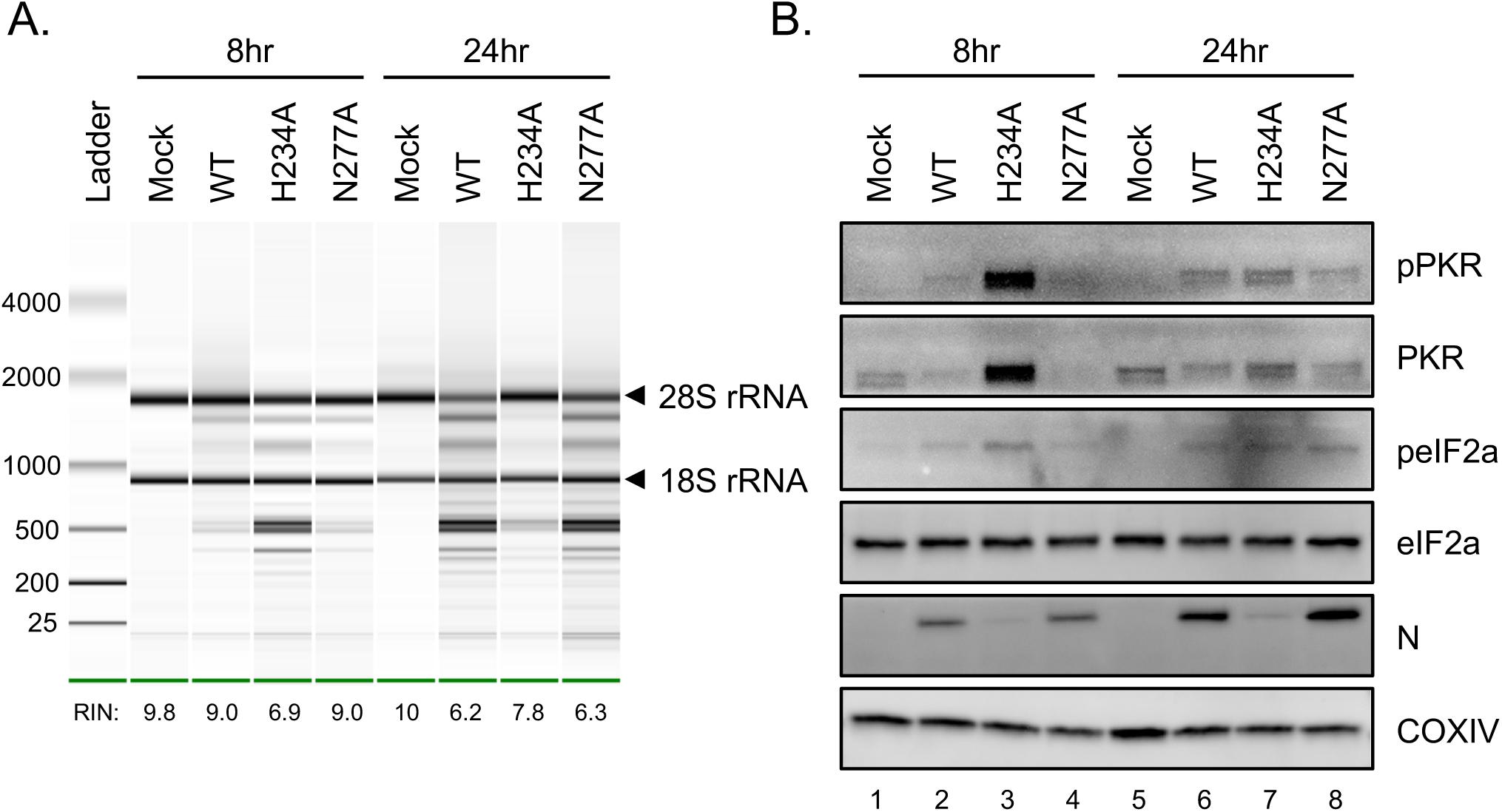
Enhanced innate immune responses during Nsp15_H234A_ SARS-CoV-2 infection. A549-ACE2 cells were uninfected or infected by Nsp15_WT_, Nsp15_H234A_, and Nsp15_N277A_ rSARS-CoV-2 at MOI of 5 for 8 and 24 h. (A) Total RNA extracted for rRNA integrity analysis by use of the Agilent 2100 bioanalyzer. (B) Cell lysates harvested for western blot analysis.

**Fig. S2.**
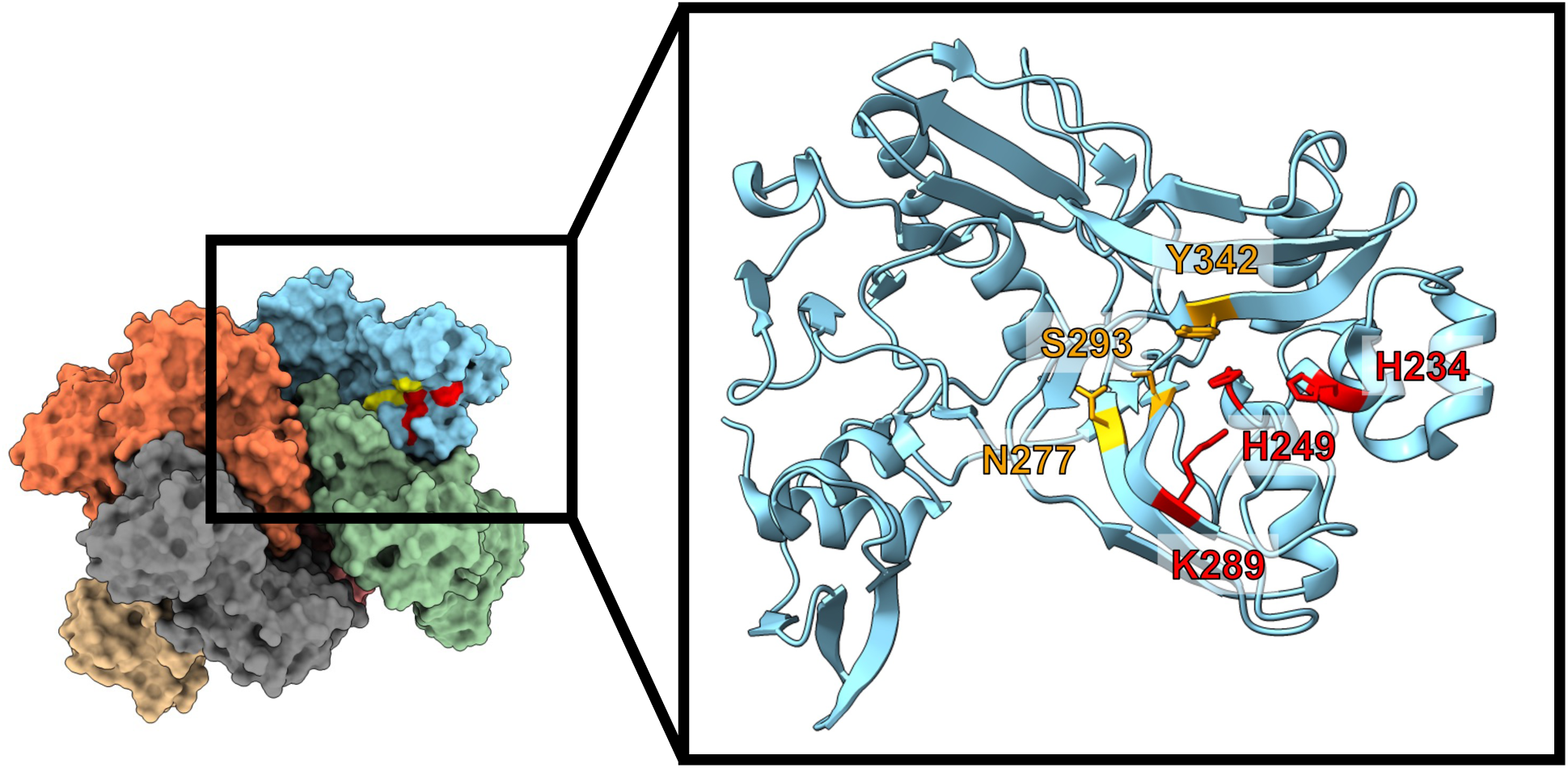
Amino acids associated with EndoU activity and uridine specificity within SARS-CoV-2 Nsp15. Cryo-EM structure of SARS-CoV-2 Nsp15 hexamer (PDB ID: 7K0R) is shown with the catalytic triad residues (H234, H249, K289) labeled in red, and residues involved in uridine discrimination (N277, S293, Y342) colored yellow. Images were rendered in ChimeraX v1.8.

**Fig. S3.**
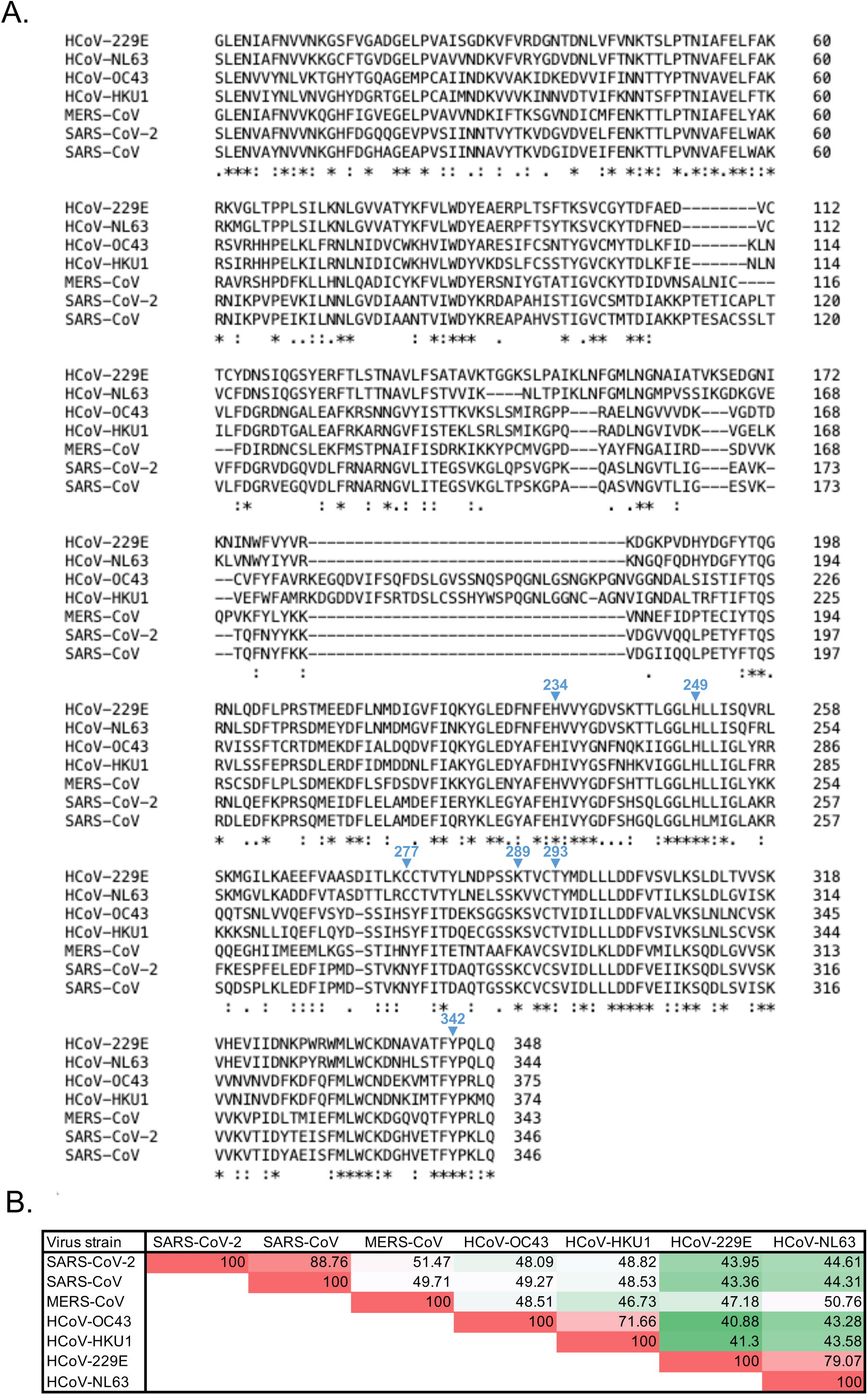
Sequence identity of Nsp15 across human coronaviruses. Nsp15 sequences from SARS-CoV-2 (Wuhan-Hu-1 strain, GenBank: NC045512), SARS-CoV (Urbani strain, GenBank: AY278741), MERS-CoV (EMC/2012 strain, GenBank: NC019843), HCoV-OC43 (ATCC VR-759, GenBank: AY585228), HCoV-HKU1 (GenBank: NC006577), HCoV-229E (ATCC VR-740, GenBank: AF304460), HCoV-NL63 (Amsterdam 1 strain, GenBank: NC005831) were acquired from the Nucleotide database, NCBI. Asterisk (*) = fully conserved; colon (:) = conserved positions containing residues with strongly similar properties; period (.) = conserved positions containing residues with weakly similar properties. (A) The amino acid sequences from HCoV Nsp15 aligned by Clustal 2.1. (B) The percentage of sequence identity among HCoV Nsp15.

